# Genome-based metabolic and phylogenomic analysis of the type strains of *Terrisporobacte*r *mayombei*, *Terrisporobacter petrolearius* and *Terrisporobacter glycolicus*

**DOI:** 10.1101/2023.08.02.551728

**Authors:** Tim Böer, Frank R. Bengelsdorf, Mechthild Bömeke, Rolf Daniel, Anja Poehlein

## Abstract

To date, three validly published type strains of the genus *Terrisporobacter* are covered by draft genome sequences, and the genes and pathway responsible for acetogenesis have not been analyzed. Here, we report complete genome sequences of the bacterial type strains *Terrisporobacter petrolearius* JCM 19845^T^, *Terrisporobacter mayombei* DSM 6539^T^ and *Terrisporobacter glycolicus* DSM 1288^T^. Functional annotation, KEGG pathway module reconstructions and screening for virulence factors were performed. Various species-specific vitamin, cofactor and amino acid auxotrophies were identified and a model for acetogenesis of *Terrisporobacter* was constructed. The complete genomes harbored a gene cluster for the reductive proline-dependent branch of the Stickland reaction located on an approximately 21 kb plasmid, which is exclusively found in the *Terrisporobacter* genus. Phylogenomic analysis of available *Terrisporobacter* genomes suggested a reclassification of most isolates as *T. glycolicus* into *T. petrolearius*.

## Introduction

Acetogenic bacteria are of high interest to biotechnology as they enable the conversion of industrial waste gases to valuable biocommodities by synthesis gas fermentation or microbial electrosynthesis (MES). These carbon-recycling technologies can contribute to the current efforts towards achieving climate neutrality. The genus *Terrisporobacter* comprises isolates capable of performing MES and autotrophic growth on H_2_ and CO_2_ through acetogenesis [1–3]. *Terrisporobacter* consists of mesophilic, Gram-positive and strictly anaerobic members. The cells appear as slightly curved rods, form endospores and are motile by flagella [4]. The four validly published type strains are *Terrisporobacter mayombei* DSM 6539^T^, *Terrisporobacter glycolicus* DSM 1288^T^, *Terrisporobacter petrolearius* JCM 19845^T^ and *Terrisporobacter hibernicus* NCTC 14625^T^. The ability to utilize H_2_ and CO_2_ via the Wood-Ljungdahl pathway (WLP) has only been shown for *T. mayombei* (Kane et al. 1991). Acetogenesis has also been described for the isolates *T. glycolicus* DSM 13865 and ATCC 29797, and genes required for the Wood-Ljungdahl pathway are present in all available *Terrisporobacter* draft genomes [2,5]. So far, no detailed genome-based analysis is available for the genes and pathways involved in acetogenic growth of *Terrisporobacter* strains. The first *Terrisporobacter* species discovered was *T. glycolicus* DSM 1288^T^ (formerly *Clostridium glycolicum*), which was isolated from mud of a stagnant pond near the Chesapeake and Ohio canal (USA) in 1962 [6]. *T. mayombei* DSM 6539^T^ (formerly *Clostridium mayombei*) was isolated as the second *Terrisporobacter* type strain from gut contents of the soil-feeding termite *Cubitermes speciosus* in 1991 [3]. The genus *Terrisporobacter* was proposed in 2014 and *T. glycolicus* DSM 1288^T^ was validly described as a type species [4]. The third type strain *T. petrolearius* JCM 19845^T^ was isolated in 2015 using a petroleum reservoir sample from an oilfield in Shengli (China) [7]. *Terrisporobacter othiniensis* (08-306576) and *Terrisporobacter muris* (DSM 29186) were described as different strains. However, both strains appeared to be the same species, but these names have not been validly published [8,9]. In 2023, *Terrisporobacter hibernicus* (NCTC 14625^T^) was validly published as a fourth type strain [10]. *Terrisporobacter* members have been found to be strongly associated to pig manure and in particular *T. glycolicus* DSM 1288^T^ has been discussed as potential emerging anaerobic pathogen [11,12]. In addition, some members show a positive correlation with the generation of bioelectricity in microbial fuel cells, indicating their electricigenic potential [13]. Correspondingly, *Terrisporobacter* strains have been found at the cathode of MES cells [1]. These insights support the use of strains from the genus *Terrisporobacter* as biocatalysts in biotechnology. The availability of high-quality genome sequences is a prerequisite for further metabolic investigations. Nevertheless, high-quality genome sequences were not available for three published type strains of the *Terrisporobacter* genus. We therefore sequenced and assembled the genomes of *T. petrolearius* JCM 19845^T^, *T. mayombei* DSM 6539^T^ and *T. glycolicus* DSM 1288^T^ to perform a genome-based metabolic and phylogenomic analysis of these *Terrisporobacter* type strains.

## Materials and Methods

### Cultivation and DNA Extraction

Genomic DNA was extracted from cells of a *T. petrolearius* JCM 19845^T^ culture derived from the Microbe Division of the Japan Collection of Microorganisms (JCM; Tsukuba; Japan). *T. glycolicus* DSM 1288^T^ and *T. mayombei* DSM 6539^T^ were grown from lyophilised cultures derived from the German Collection of Microorganisms and Cell Cultures (DSMZ; Braunschweig, Germany) in an overnight culture, incubated at 30 °C in DSM 311 medium supplemented with glucose (30 mM). Cell pellets were obtained by centrifugation of 10 ml culture at 13,000 g for 5 min. DNA was isolated by using the instructions for cell samples of the MasterPure™ Complete DNA and RNA purification kit (Epicentre, Madison, WI, USA).

### Genome Sequencing, Assembly and Annotation

Illumina sequencing libraries were prepared using the Nextera XT DNA sample preparation kit (Illumina, San Diego, CA, USA) and sequenced with a MiSeq system and v3 chemistry (600 cycles) following the instructions of the manufacturer (Illumina, San Diego, CA, USA). Sequencing kits and program versions for Nanopore sequencing are shown in the Supplementary Table S1. The Nanopore sequencing libraries were prepared with 1.5 µg high-molecular-weight DNA using the ligation sequencing kit 1D and the native barcode expansion kit as recommended by the manufacturer (Oxford Nanopore Technologies, Oxford, UK). Nanopore sequencing was conducted for 72 h with the MinION device Mk1B, the SpotON flow cell R9.4.1 and the MinKNOW software following the instructions of the manufacturer (Oxford Nanopore Technologies, Oxford, UK) Demultiplexing and base calling of Nanopore sequencing data were performed with the Albacore/Guppy software. Quality control and adapter trimming of paired-end Illumina sequences was performed with FastQC (v0.11.9) and Trimmomatic (v0.39; LEADING: 3, TRAILING: 3, SLIDINGWINDOW:4:15, MINLEN:50) [14]. Porechop (v0.2.4) was used for adapter trimming of Nanopore reads, which were subsequently filtered to a minimal read length of 5 kb [15]. Trimmed paired-end Illumina and filtered long Nanopore reads were used as input of Unicycler (v0.5.0) for a *de novo* hybrid genome assembly [16]. The resulting assembly graph was inspected with Bandage (v0.8.1) [17]. Annotation of the resulting genomes was performed with Prokka (v1.14.6) [18] and the locus tag TEMA for *T. mayombei* DSM 6539^T^, TEPE for *T. petrolearius* JCM 19845^T^ and TEGL for *T. glycolicus* DSM 1288^T^ were chosen. BUSCO scores were obtained using BUSCO (v5.4.4) with the lineage dataset clostridia_odb10 and the mode for prokaryotic genomes [19].

### Genome-based metabolic and phylogenomic analysis

Functional annotation of genes to KEGG categories and the reconstruction of KEGG pathway modules was conducted with BlastKOALA [20]. For metabolites occurring with multiple KEGG pathway modules only the most complete ones were chosen for visualization. Assignment of genes into cluster of orthologous genes (COG) and subsequent functional annotation into COG categories was performed with the eggNOG mapper (v2.1.9) using the eggNOG 5.0 database [21,22]. Barcharts and heatmaps were plotted in RStudio (v2022.06.0) with the ggplot2 package (v3.4.1) [23,24]. Pan/core genome analysis was conducted with OrthoVenn2 clustering genes, identifying orthologous clusters and assigning Gene Ontology (GO) annotations [25]. The identification and classification of insertion sequence (IS) elements was performed with the ISESCAN pipeline (v1.7.2.3) [26]. Orthologous genes for the Wood-Ljundgdahl pathway and the Stickland reactions were identified by using proteinortho (v6.0.31) running with default parameters [27]. Gene cluster visualizations were prepared with clinker (v1.32) [28]. Virulence factor screening was performed with the VFanalyzer pipeline using the Virulence Factor Database (VFDB) [29]. The phylogenomic analysis was performed with all *Terrisporobacter* genomes available at NCBI Genbank using pyani (v0.2.12) and the MUMmer alignment option [30]. ANIm percentage identity and alignment length values are shown in S2 Table. The dDDH values for *T. petrolearius* were calculated by the Type (Strain) Genome Server (TYGS) [31].

## Results and Discussion

### Genome Assembly and Annotation

Details for the genome sequencing, assembly and annotation of the three *Terrisporobacter* type strains are summarized in Table 1. Each genome harbors a circular chromosome ranging from 4.03 - 4.09 Mb, and an additional circular plasmid ranging from 21.29 – 21.89 Kb (pTM1, pTP1 and pTG1). An additional circular plasmid was obtained for *T. mayombei* DSM 6539^T^ (pTM2, 8.20 Kb) and *T. petrolearius* JCM 19845^T^ (pTP2, 6.74 Kb). The GC content of the chromosomes varied between 28.7 – 28.9 %. Prokka annotation resulted in 4,165, 4,208 and 4,038 potential genes for the genomes of *T. mayombei, T. petrolearius* and *T. glycolicus,* respectively. All three genomes contained 12 genes coding for 5S, 16S and 23S rRNA. One CRISPR repeat region was detected in the genomes of *T. petrolearius* and *T. glycolicus* with 67 and 28 spacers, respectively. The genome of *T. mayombei* contained two CRISPR repeat regions with 12 and 25 spacers. BUSCO scored a 100 % completeness for all three genomes with 98.8 % single copy (S) and 1.2 % duplicated (D) BUSCO marker genes. No fragmented (F) or missing (M) BUSCO marker genes were identified.

**Table 1.** Genome features of the three *Terrisporobacter* type strains *T. petrolearius* JCM 19845^T^, *T. mayombei* DSM 6539^T^, and *T. glycolicus* DSM 1288^T^.

|  | Genome |  |  |
| --- | --- | --- | --- |
| Feature | *T. mayombe*<br>DSM 6539<sup>T</sup> | *T. petrolearius*<br>JCM 19845<sup>T</sup> | *T. glycolicus*<br>DSM 1288<sup>T</sup> |
| Chromosome size<br>(bp) | 4,034,168 | 4,096,312 | 4,039,277 |
| Number of Illumina<br>reads (250 bp) | 1,495,093 | 1,218,094 | 4,189,121 |
| Number of Nanopore<br>reads/mean length<br>(bp) | 207,411/11,100 | 295,055/6,700 | 113,279/20,700 |
| Chromosome mean<br>coverage<br>(Illumina/Nanopore) | 116/557 | 122/465 | 444/552 |
| Plasmid size (bp) | pTM1: 21,885;<br>pTM2: 8,218 | pTP1: 21,288;<br>pTP2: 6,738 | pTG1: 21,319 |
| Plasmid mean<br>coverage<br>(Illumina/Nanopore) | pTM1: 417/2,314<br>pTM2: 1,008/3,389 | pTP1: 557/2,416<br>pTP2: 428/2,161 | pTG1: 1,526/1,573 |
| GC content (%) | 28.8 | 28.9 | 28.7 |
| Genes | 4,165 | 4,203 | 4,038 |
| CDS | 3,915 | 3,932 | 3,826 |
| Functional proteins | 2,033 | 2,112 | 2,114 |
| Hypothetical proteins | 1,882 | 1,820 | 1,712 |
| rRNAs<br>(5S; 16S; 23S) | 12; 12; 12 | 12; 12; 12 | 12; 12; 12 |
| tRNAs | 94 | 93 | 93 |
| tmRNA | 1 | 1 | 1 |
| ncRNAs | 119 | 141 | 82 |
| CRISPR repeats | 2 | 1 | 1 |
| BUSCO scores | C:100.0 %<br>[S:98.8 %, D:1.2 %]<br>F:0.0%, M:0.0 %<br>n:247 | C:100.0 %<br>[S:98.8 %, D:1.2 %]<br>F:0.0 %, M:0.0 %<br>n:247 | C:100.0 %<br>[S:98.8 %, D:1.2 %]<br>F:0.0 %, M:0.0 %<br>n:247 |

### Functional annotation and pan/core genome analysis

The functional annotation into COG categories in the *Terrisporobacter* type strain genomes is visualized in Fig 1A. The eggNOG-mapper assigned categories to 86.9 % of coding sequences for *T. mayombei*, 88.1 % of coding sequences for *T. petrolearius* and 89 % for *T. glycolicus*. The genes of the three *Terrisporobacter* type strain genomes distributed uniformly across the different COG categories, with the exception of the category replication, recombination and repair (L). With 258, the *T. petrolearius* contained significantly more genes than the other two type strains of *T. mayombei* (161) and *T. glycolicus* (150). The pan genome of the three genomes comprised 3,548 gene clusters of which 2,930 gene clusters formed the core genome (Fig 1B). The genome of *T. mayombei* and *T. glycolicus* each harbored 15 exclusive gene clusters, while the genome of *T. petrolearius* harbored 38 exclusive gene clusters. *T. mayombei* contained four exclusive gene clusters, including gene clusters for acyltransferase activity (GO:0016746), phosphorelay sensor kinase activity (GO:0000155), DNA binding (GO:0003677) and metal ion binding (GO:0046872). Furthermore, *T. mayombei* possessed two exclusive gene clusters with GO annotations for the regulation of DNA-templated transcription (GO:0006355) and cobalamin biosynthesis (GO:0009236). *T. petrolearius* contained ten exclusive gene clusters with GO annotations of molecular function, comprising gene clusters for ATP-binding (GO:0005524), phosphorelay response regulator activity (GO:0000156), metal ion binding (GO:0046872), spermidine-importing ATPase activity (GO:0015595), enterobactin transmembrane transporter activity (GO:0042931), transmembrane transporter activity (GO:0022857), methionyl-tRNA formyltransferase activity (GO:0004479), sequence-specific DNA binding (GO:0043565) and two gene clusters encoding ATP hydrolysis activity (GO:0016887). Four exclusive gene clusters with over 80 predicted proteins in total were annotated for transposition (GO:0032196) and DNA-mediated transposition (GO:0006313). In addition, exclusive gene clusters were identified with GO annotations for osmosensory signaling via phosphorelay pathway (GO:0007234), shikimate metabolic process (GO:0019632), antibiotic biosynthetic process (GO:0017000), regulation of DNA-templated transcription (GO: 0006355), DNA integration (GO:0015074) and N-terminal protein amino acid acetylation (GO:0006474). *T. glycolicus* contained two exclusive gene clusters with annotated molecular function, including gene clusters for nickel cation transmembrane transporter activity (GO:0015099) and metal ion binding (GO:0046872). Other assigned GO annotations of exclusive gene clusters comprised regulation of systemic acquired resistance (GO:0010112), transmembrane transport (GO:005505085), ethanolamine catabolic process (GO:00463336), regulation of DNA- templated transcription (GO: 0006355), polysaccharide biosynthetic process (GO:0000271), nickel cation transport (GO:0015675) and teichoic acid biosynthetic process (GO:0019350). Remaining exclusive gene clusters for the three *Terrisporobacter* genomes did not receive functional annotations.

**Fig 1.**
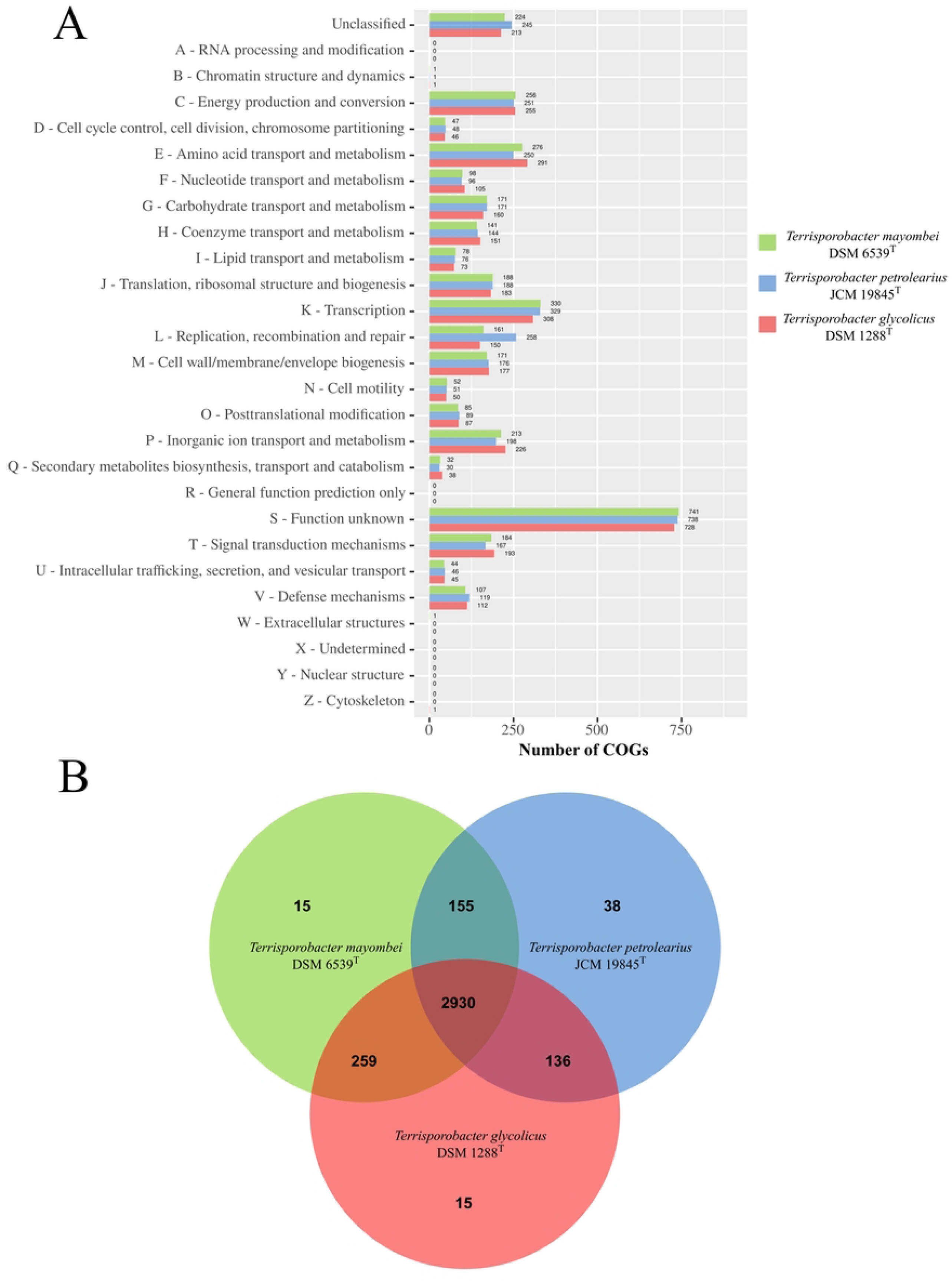
Functional annotation and pan/core genome analysis. A. Functional annotation of the *Terrisporobacter* type strain genomes into COG categories. The y-axis shows the different COG categories and the x-axis the number of genes assigned to each category by eggNOG-mapper. The bar color indicates the number of genes for the three *Terrisporobacter* type strains *T. mayombei* (green), *T. petrolearius* (blue) and *T. glycolicus* (red). **B** Pan/core genome analysis of three *Terrisporobacter* type strains. The Venn diagram visualizes the exclusive and shared gene clusters for *T. mayombei* (green), *T. petrolearius* (blue) and *T. glycolicus* (red) created with OrthoVenn2.

KEGG pathway analysis assigned 49.9 % of genes for *T. mayombei*, 51.3 % of genes for *T. petrolearius* and 52.4 % of genes for *T. glycolicus* into functional categories (Fig 2). Between the analyzed three *Terrisporobacter* type strains genes distributed uniformly to the different functional categories, with the exception of the category unclassified: genetic information processing. In this category, 113 genes were assigned from the genome of *T. petrolearius*, while only 32 genes derived from the genome of *T. mayombei* and 28 genes from the genome of *T. glycolicus*.

**Fig 2.**
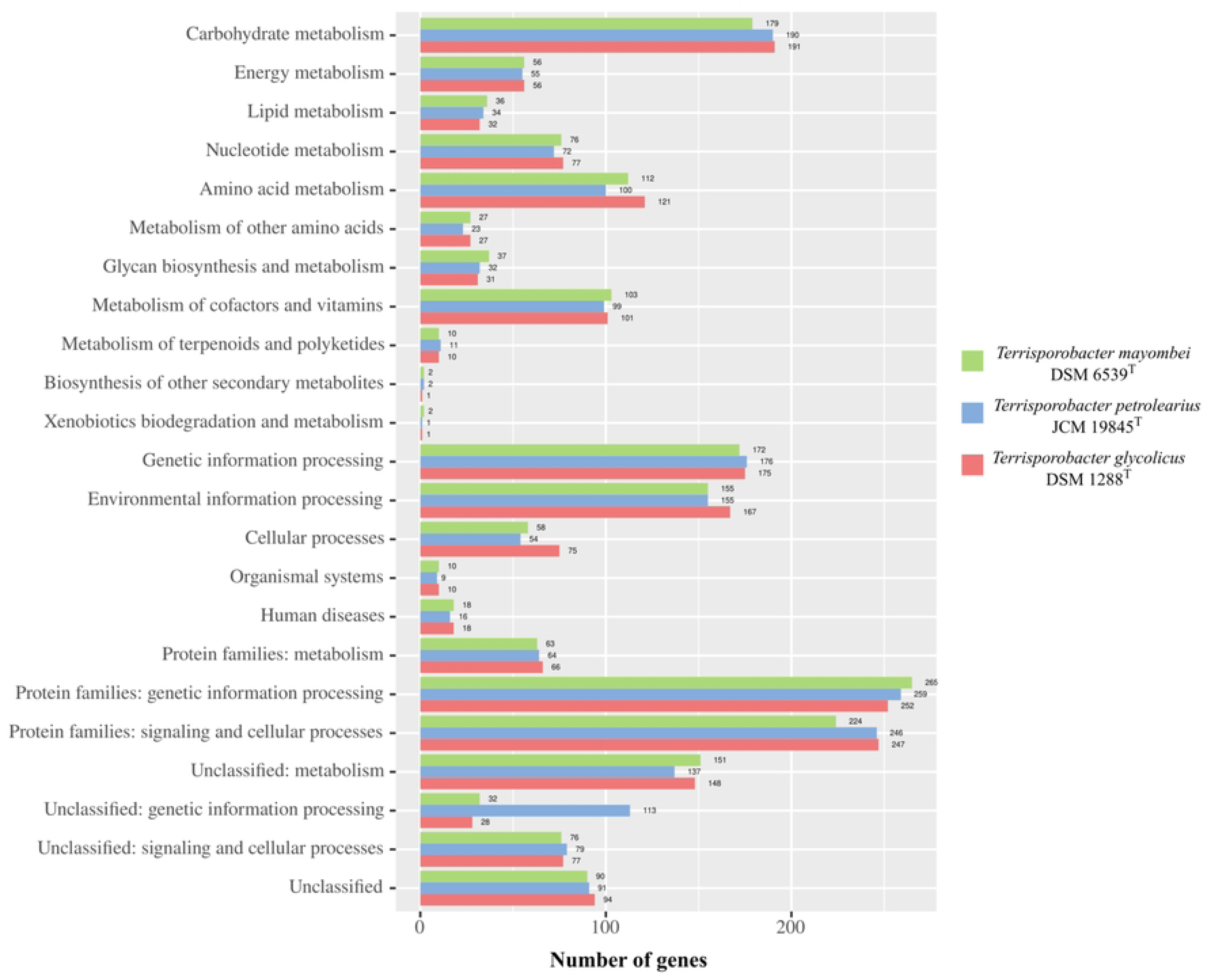
Functional annotation of genes from the *Terrisporobacter* type strain genomes into KEGG categories. The y-axis shows the different functional KEGG categories and the x-axis the number of genes assigned to each category by BlastKOALA. The bar color indicates the number of genes assigned for the three *Terrisporobacter* type strains *T. mayombei* (green), *T. petrolearius* (blue) and *T. glycolicus* (red).

The functional annotations into COG and KEGG categories match with the described *Terrisporobacter* substrate spectrum specializing on the fermentation of carbohydrates and amino acids [3,6,7]. In both functional annotation approaches the gene distribution to the different categories was very similar between the three *Terrisporobacter* type strain genomes with the only exception of the increased gene number in *T. petrolearius* assigned to the categories of unclassified: genetic information processing (KEGG) or replication, recombination and repair (COG). This difference was also identified by pan/core genome analysis, with the four exclusive gene clusters encoded by *T. petrolearius* containing over 80 predicted proteins for transposition. In order to investigate the difference in transposase abundance the presence of IS elements in the genomes were analyzed with the ISESCAN pipeline (Fig 3). In total 94 IS elements were identified for *T. petrolearius*, while only four and six IS elements were detected in the *T. mayombei* and *T. glycolicus* genomes, respectively. All three genomes harbored IS elements from the IS21 family. One element of family IS66 was present in the genomes of *T. petrolearius* and *T. glycolicus*. The family IS607 was only detected with one element in *T. glycolicus*. The families IS3, IS110 and IS256 were only recorded in *T. petrolearius* and could be identified as the major source of the detected difference in transposase abundance between the three analyzed *Terrisporobacter* genomes. The families IS3 and IS256 contain transposases with the common acidic triad of Asp, Asp and Glu at the active site (DDE transposases). The IS110 family characteristically comprise of IS elements with transposases containing an Asp, Glu, Asp, Asp motif at the active site (DEDD transposase). These IS elements differ from classical DDE transposases as they do not contain terminal inverted repeats and upon integration do not produce flanking target DNA repeats [32].The identified differences indicate an increased genome plasticity of *T. petrolearius* in comparison to the other two type strains [33].

**Fig 3.**
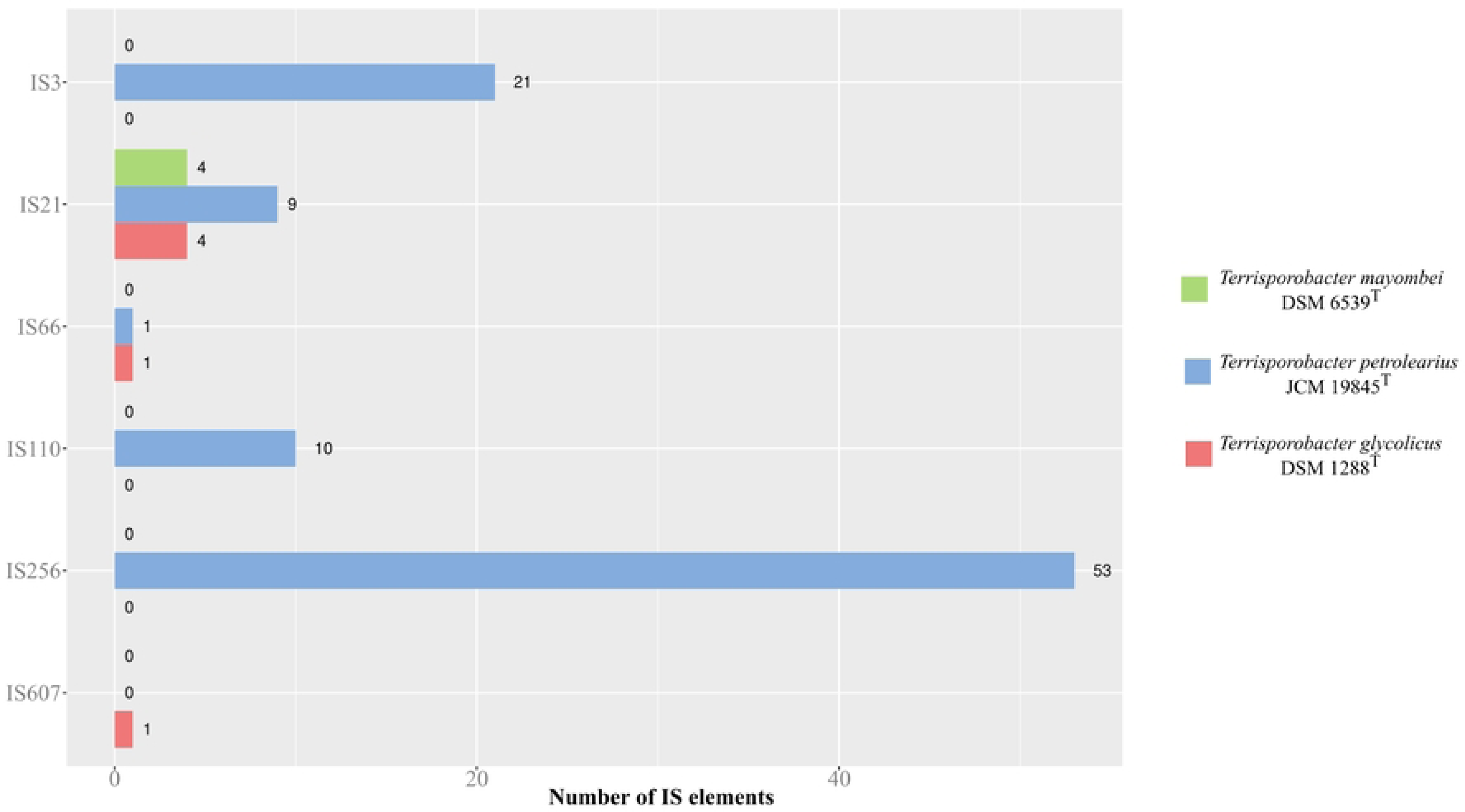
Distribution of IS elements and classification into IS families. The y-axis shows the different IS families and the x-axis the number of IS elements assigned to each IS family by ISESCAN. The bar color indicates the number of genes assigned for the three *Terrisporobacter* type strains *T. mayombei* (green), *T. petrolearius* (blue) and *T. glycolicus* (red).

For a more detailed metabolic investigation of the three *Terrisporobacter* type strain genomes, we analyzed and compared regarding the pathway modules of different KEGG categories (Fig 4). Within the category carbohydrate metabolism, the modules for glycolysis, pyruvate oxidation, pentose phosphate pathway, phosphoribosyl diphosphate (PRPP) biosynthesis, glycogen biosynthesis, nucleotide sugar biosynthesis from glucose and UDP-N-acetyl-D-glucosamine biosynthesis were complete in all three genomes. The modules for gluconeogenesis, citrate cycle, semi-phosphorylative Entner-Doudoroff pathway, D-galacturonate degradation, galactose degradation, glycogen degradation, nucleotide sugar biosynthesis from galactose and propanoyl- CoA metabolism were incomplete in every genome. The module for gluconeogenesis was identified as incomplete due to the lack of a phosphoenolpyruvate (PEP) carboxykinase (both ATP- and GTP-dependent) catalyzing the conversion of oxaloacetate into PEP. Nevertheless, gluconeogenesis remains functional as acetyl-CoA is carboxylated to pyruvate by a pyruvate ferredoxin oxidoreductase. This reaction links the Wood-Ljungdahl pathway to gluconeogenesis and the incomplete citrate cycle, thereby allowing the generation of building blocks for biosynthesis during autotrophic growth [34]. With an incomplete traditional, semi- and non- phosphorylative Entner-Doudoroff pathway, hexoses are metabolized only via glycolysis. Fermentation of pentoses such as xylose can occur via the non-oxidative pentose phosphate pathway and has been described for the type strains of *T. mayombei* and *T. glycolicus* [3,6]. All three analyzed genomes contain an incomplete citrate cycle and are not able to degrade glycogen and galactose. In the category metabolism of cofactors and vitamins every genome contained complete gene sets for the modules of thiamine salvage, NAD biosynthesis, coenzyme A biosynthesis, C1-unit interconversion and the two cobalamin biosynthesis modules. The module for the biosynthesis of pyridoxal-P was only complete in *T. mayombei* and *T. glycolicus*, while genes for this module were not present in *T. petrolearius*. Both modules for the biosynthesis of biotin were only complete in *T. mayombei* and incomplete in the other two type strains. The modules for the biosynthesis of thiamine, riboflavin, pantothenate, pimeloyl-ACP, lipoic acid, tetrahydrofolate, tetrahydrobiopterin, L-threo-tetrahydrobiopterin, molybdenum cofactor, siroheme, heme and tocopherol/tocotorienol were incomplete in all three *Terrisporobacter* type strains. Siroheme biosynthesis was incomplete as all three genomes lack the sirohydrochlorin ferrochelatase (sirB). The genome of *T. petrolearius* is additionally missing genes encoding the glutamyl-tRNA reductase (hemA) and the glutamate-1-semialdehyde 2,1-aminomutase (hemL). However, all three genomes encode sulfite reductases requiring siroheme as essential prosthetic group [35]. The incomplete module for heme biosynthesis in all three genomes imply the dependence on heme acquisition from the environment or the host during potential pathogenesis [36]. The category amino acid metabolism contained complete gene sets for the modules threonine biosynthesis, cysteine biosynthesis, lysine biosynthesis, polyamine biosynthesis and histidine degradation in all three genomes. The modules for tyrosine and phenylalanine biosynthesis were only complete for *T. mayombei* and *T. glycolicus*, while the module for histidine biosynthesis was only complete in *T. petrolearius* and *T. glycolicus*. Furthermore, the modules for leucine and tryptophan biosynthesis were only complete in *T. glycolicus*. The modules for the biosynthesis of serine, methionine, valine, isoleucine, arginine, proline, gamma-aminobutyric acid GABA and the shikimate pathway were incomplete in all three *Terrisporobacter* type strains. This also applied to the modules for the urea cycle and GABA shunt. The identified incomplete modules in the categories metabolism of cofactors and vitamins and amino acid metabolism explain the observed dependence on the addition of vitamins and amino acids for significant growth of *Terrisporobacter* members [3,6,7]. The reconstruction of KEGG pathway modules showed auxotrophies for different vitamins/cofactors and amino acids in all three genomes. Furthermore, several species-specific vitamin and amino acid auxotrophies were identified. In the category energy metabolism complete modules were detected in every *Terrisporobacter* type strain for reductive acetyl-CoA pathway, phosphate acetyltransferase-acetate kinase pathway and F-type ATPase. The modules reductive pentose phosphate cycle, formaldehyde assimilation and assimilatory nitrate reduction were incomplete in all three *Terrisporobacter* strains. No complete modules were present for the category of biosynthesis of terpenoids and polyketides. The signature modules for beta-lactam resistance and acetogen were complete in all three *Terrisporobacter* type strains. The prerequisites for growth on H_2_ and CO_2_ through acetogenesis are fulfilled for every *Terrisporobacter* type strain analyzed, as shown by the complete modules for the reductive acetyl-CoA pathway (WLP), phosphate acetyltransferase-acetate kinase pathway and the complete signature module for acetogens.

**Fig 4.**
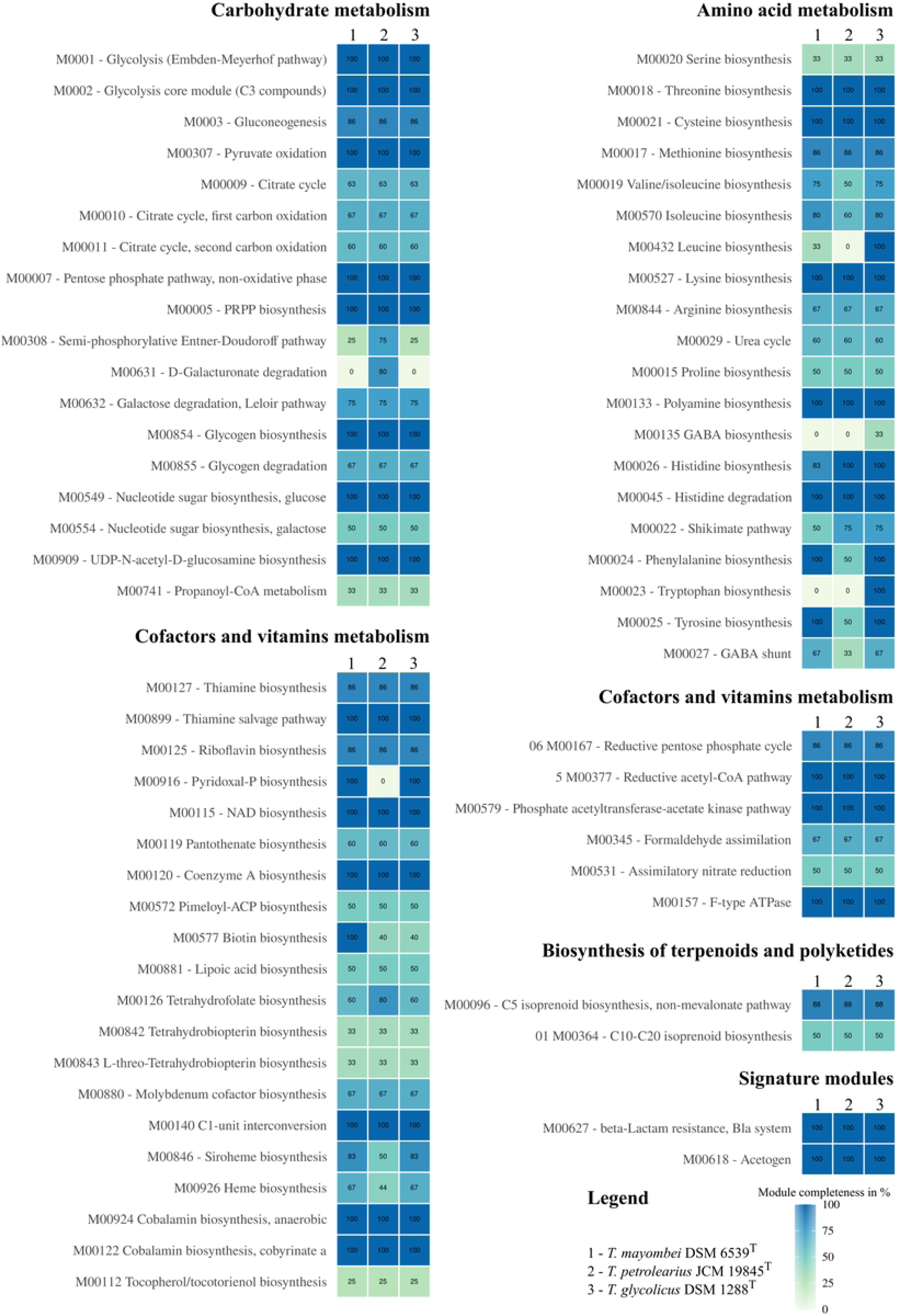
Completeness of KEGG pathway modules for KEGG categories. The completeness of KEGG modules is given in percentage from the total amount of required blocks and the amount of detected blocks in a pathway module for the three *Terrisporobacter* type strains *T. mayombei* (1), *T. petrolearius* (2) and *T. glycolicus* (3). For metabolites with multiple KEGG modules only the most complete modules detected are shown. The legend for the color gradient is shown in the bottom right.

### Pathway reconstruction for acetogenesis

The genes necessary to employ the WLP were present in all three type strains of the *Terrisporobacter* genus. The detected genes for the WLP are shown in Fig 5A with the corresponding locus tags for each genome. Overall, the genomic organization of the WLP genes in the *Terrisporobacter* type strains resembled the structure detected in the acetogen *Clostridioides difficile* [37]. The methyl-branch starts with the reversible hydrogen-dependent CO_2_ reductase complex (HDCR) using H_2_ as a direct electron donor for the reduction of CO_2_ to formate by a formate dehydrogenase (FDH). A potential HDCR-complex was encoded by a gene cluster consisting of a selenocysteine-containing formate dehydrogenase H (*fdhF*), two copies of the hydrogenase-4 component A (*hyfA*), a NADP-reducing hydrogenase subunit (*hndD*), a putative formate transporter (*focA*) and a sulfur carrier protein (*fdhD*). In *C. difficile* the HDCR-complex gene cluster did not contain a formate transporter and *fdhD* was located adjacent to the *fdhF* gene, while in *Terrisporobacter* it was located at the end of the gene cluster (Fig 5B). In addition to the HDCR-complex, the oxidation of H_2_ in *Terrisporobacter* can be performed by orthologs of the electron-bifurcating hydrogenase *HydABC* (TEMA_26770-26790, TEPE_39410-39430, TEGL_37760-37780). The activation of formate with tetrahydrofolate (THF) is carried out under consumption of one ATP by the formyl-THF synthetase (FTS) producing N^10^-formyl-THF, which is subsequently transformed to N^5^,N^10^-methenyl-THF via cyclization and dehydration by the methenyl-THF cyclohydrolase (MTC). The methylene-THF dehydrogenase (MTD) reduces N^5^,N^10^-methenyl-THF to N^5^,N^10^-methylene-THF, which is then further reduced to N^5^-methyl-THF by the methylene-THF reductase (MTR). Finally, the methyl-group is transferred to a corrinoid- iron-sulfur protein (CoFeSP) by the methyl-transferase (MT). In the carbonyl-branch the carbon monoxide dehydrogenase (CODH) reduces CO_2_ to a carbonyl-group using reduced ferredoxin. This carbonyl group is fused with the methyl-group from the methyl-branch and the cofactor coenzyme A (CoA), which is catalyzed by the acetyl-CoA synthetase (ACS) and releases CoFeSP. Acetyl-CoA is converted to acetyl phosphate (Acetyl-P) by a phosphotransacetylase (PTA) or alternatively by a phosphotransbutyrylase (PTB). The transfer of the phosphate group to ADP forming ATP is catalyzed by an acetate or alternatively butyrate kinase (ACK*/*BUK) and leads to the formation of the final product acetate. The genus *Terrisporobacter* is reported to produce only acetate as a fermentation product of the WLP (homoacetogenesis) [2]. The majority of genes involved in the WLP were organized as a gene cluster of approximately 18 kb. These gene clusters showed the same organization as in *C. difficile* or acetogenic bacteria belonging to the genus *Clostridium* (Fig 5C) [38]. Energy conservation is accomplished with the RNF-complex (*RnfABCDEG*; TEMA_15820-15870, TEPE_33980-33930, TEGL_15820-15870) by coupling the electron transfer from reduced ferredoxin to NAD with the translocation of protons or sodium ions across the cytoplasmic membrane. A membrane-bound ATP synthase can then use the created ion gradient to produce ATP [39]. The redox pool of NADH and reduced ferredoxin is linked to the cellular redox pool (NADPH) with a *Sporomusa* type Nfn (STN) electron-bifurcating transhydrogenase (*StnABC*, TEMA_09310-09330, TEPE_23120-23140, TEGL_20000-20020) [40].

**Fig 5.**
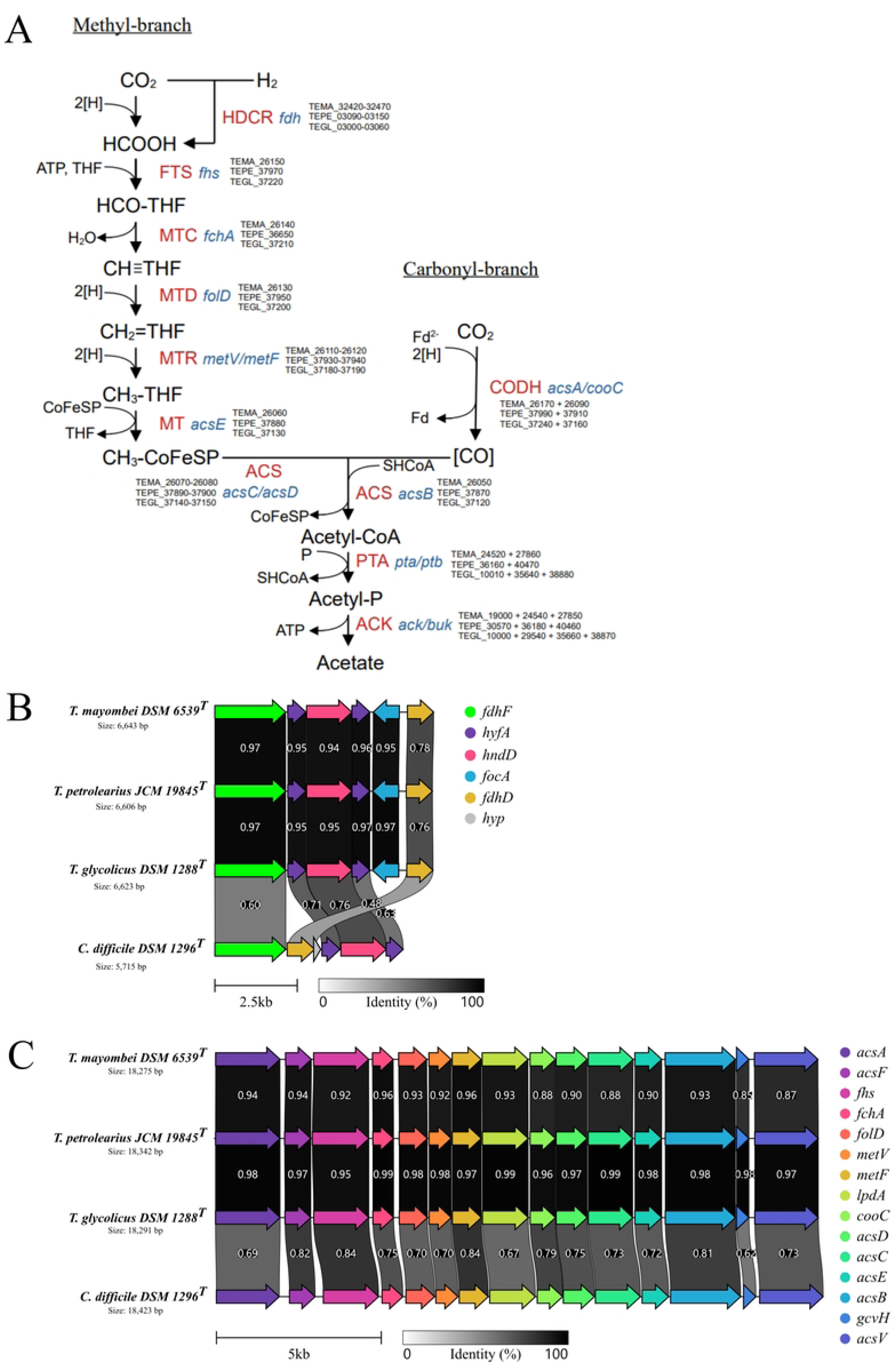
**Reconstruction of the Wood-Ljungdahl pathway in *Terrisporobacter***. **A**: Genome-based reconstruction of the Wood-Ljungdahl pathway for the three type strains of the *Terrisporobacter* genus. Genome locus tags (black) of the catalyzing enzymes (red) and the required genes (blue) are shown for the three type strains of the *Terrisporobacter* genus (TEMA, *T. mayombei* DSM 6539^T^; TEPE, *T. petrolearius* JCM 19845^T^; TEGL, *T. glycolicus* DSM 1288^T^). **B**: Genomic organization of the HDCR-complex gene cluster in the three *Terrisporobacter* type strains in comparison to the type strain of *C. difficile* DSM 1269^T^. The following gene abbreviations were used: *fdhF*, formate dehydrogenase H; *hyfA*, hydrogenase-4 component A; *hndD*, NADP-reducing hydrogenase; *focA*, putative formate transporter; *fdhD*, sulfur carrier protein. **C**: Genomic organization of the Wood-Ljungdahl gene cluster in the three *Terrisporobacter* type strains in comparison to the type strain of *C. difficile* DSM 1269^T^. The following gene abbreviations were used: *acsA*, anaerobic carbon-monoxide dehydrogenase catalytic subunit; *acsF*, carbon monoxide dehydrogenase accessory protein; *fhs*, formyl THF synthetase; *fchA*, methenyl THF cyclohydrolase; *folD*, bifunctional cyclohydrolase/dehydrogenase; *metV*, methylene THF reductase C-terminal catalytic subunit; *metF*, methylene THF reductase large subunit; *lpdA*, dihydrolipoyl dehydrogenase; *cooC*, carbon monoxide dehydrogenase accessory protein; *acsD*, CoFeSP small subunit; *acsC*, CoFeSP large subunit; *acsE*, methyl THF CoFeSP methyltransferase; *acsB*, carbon monoxide dehydrogenase/acetyl-CoA synthase subunit beta; *gcvH*, glycine cleavage system H protein; *acsV*, corrinoid activation/regeneration protein. The following enzyme abbreviations were used: HDCR, hydrogen-dependent CO2 reductase; FTS, formyl-THF synthetase; MTC, methenyl-THF cyclohydrolase; MTD, methylene-THF dehydrogenase; MTR, methylene-THF reductase; MT, methyl-transferase; CODH, carbon monoxide dehydrogenase; ACS, acetyl-CoA synthetase; PTA, phosphotransacetylase, ACK, acetate kinase. The legend for size and sequence identity of translated genes are shown at the bottom of **B** and **C** in kilobases (kb) and from 0 % (white) to 100 % (black).

### Terrisporobacter plasmids

All three *Terrisporobacter* type strain genomes harbored a circular plasmid with an approximate size of 21 kb (pTM1, pTP1 and pTG1). *T. petrolearius* and *T. mayombei* contained an additional smaller circular plasmid of 6.7 kb (pTP2) and 8.2 kb (pTM2) in size. The annotation of the smaller plasmids pTP2 and pTM2 yielded only genes encoding hypothetical proteins. Hence, inferring possible functional roles of these plasmids was not possible. A circular plasmid of similar size (21.7 kb) as the 21 kb plasmids was also detected in the complete genome of *T. hibernicus* NCTC 14625^T^ [10]. These plasmids appeared to be a shared trait in the *Terrisporobacter* genus, as they were present with an almost identical genetic organization in the genomes of every *Terrisporobacter* type strain (Fig 6). All three larger plasmids contained a full set of genes required for the proline-dependent reductive reactions of the Stickland reaction [41]. The plasmids encoded the proprotein *prdA*, which is most likely posttranslationally cleaved resulting in one small protein containing the pyruvoyl-group for binding of D-proline and one larger scaffold protein. The proline reductase was completed by the gene products of *prdB* and *prdC*. As in *C. difficile,* only *prdB* encodes a selenocysteine residue (Sec), while the *prdC* gene did not encode a Sec as typically present in other amino acid-fermenting bacteria such as *Acetoanaerobium sticklandii* or *Clostridium botulinum* [42]. The use of a selenoprotein requires the cotranslational insertion of Sec guided by a complex machinery [43,44]. The Sec insertion complex was encoded by the *Terrisporobacter* plasmids by the three genes *selA*, *selB*, *selD* and the selenium-binding tRNA gene *selC*. In addition to genes for proline reduction, the plasmids contained genes for the partitioning proteins ParA and ParB, a DNA-invertase (Hin), four hypothetical proteins and a putative cysteine desulfurase (Csd). The plasmid of *T. mayombei* additionally encoded a protein annotated as cold shock protein (Csp). Genes for the reduction of glycine/sarcosine/betaine and the oxidative branch of the Stickland reaction were all encoded chromosomally. This also applied to the proline racemase (PrdF) required for the transformation of amino acids from the L to the D stereoisomer (TEMA_04260 + 37640/TEPE_08380/ TEGL_07940) and the membrane-bound protein PrdG (TEMA_37630 + 37650/TEPE_08370 + 08390/ TEGL_07930 + 07950). Orthologs of the putative DNA-binding protein PrdX and the sigma-54-dependent transcriptional activator protein PrdR were not identified.

**Fig 6.**
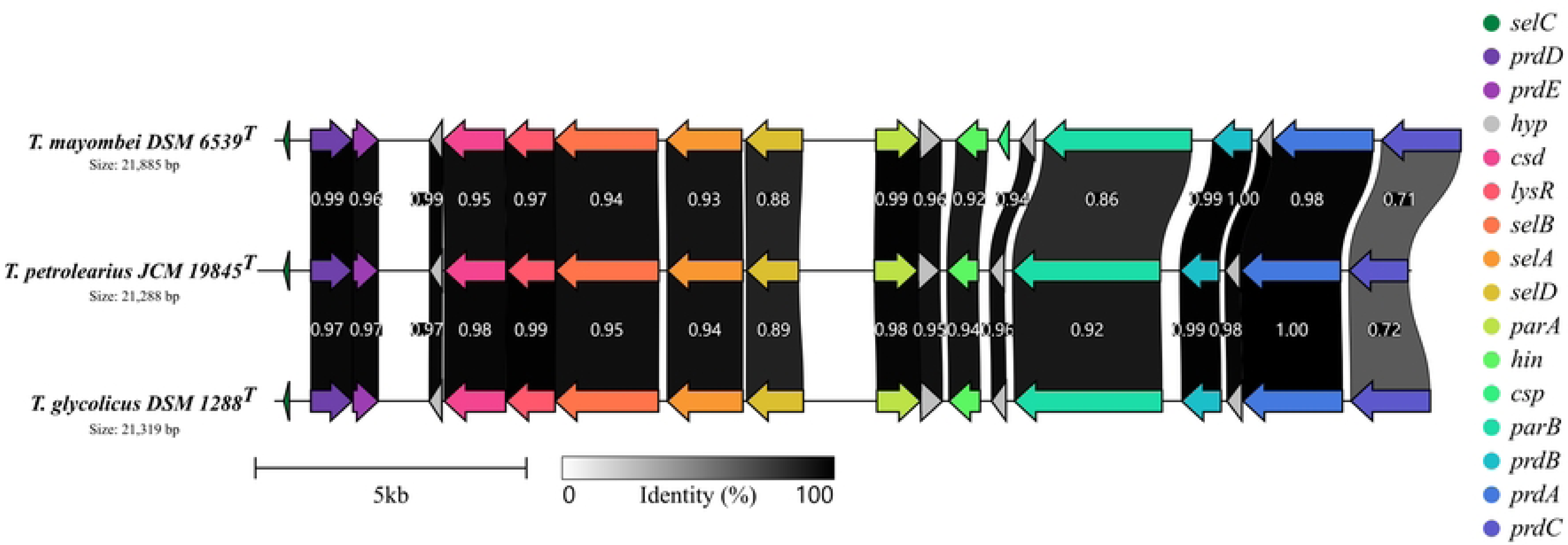
Linearized comparison of the circular plasmids in the three *Terrisporobacter* type strains. The plasmids encode a proline-dependent gene cluster for the reductive branch of the Stickland reaction. The legend for the size and the sequence identity of translated genes is shown at the bottom in kilobases (kb) and from 0 % (white) to 100 % (black). The following genes are encoded: *selC*, tRNA-Sec; *prdD*, proline reductase cluster protein prdD; *prdE*; proline reductase cluster protein prdE; *csd*, putative cysteine desulfurase; *lysR*, lysR family transcriptional regulator; *selB*, selenocysteine-specific elongation factor; *selA*, L-seryl-tRNA (Sec) selenium transferase; *selD*, selenide water dikinase; *parA*, parA family protein; *hin*, DNA-invertase; *csp*, cold shock protein; *parB*, DNA primase; *prdB*, D-proline reductase subunit gamma; *prdA*, D-proline reductase proprotein PrdA; *prdC*, proline reductase-associated electron transfer protein; *hyp*, hypothetical protein.

### Virulence factor screening

Multiple potential virulence factors were detected in the genomes of the three *Terrisporobacter* type strains comprising the VF classes adherence, regulation, toxin, antiphagocytosis, cell surface components, copper uptake, enzyme, glycosylation systems, immune evasion, iron uptake, others, secretion system, serum resistance and immune evasion and stress adaptation (Table 2). Six virulence factors for cell adherence were detected. Genes encoding a fibronectin-binding protein (*fbpA/fbp68*) and a heat shock protein (*groEL*) were found in every *Terrisporobacter* type strain. These proteins with adhesive function are characteristically present in pathogenic Gram-positive bacteria such as *C. difficile* or *Clostridium perfringens* [45–47]. Furthermore, the *Terrisporobacter* type strain genomes encoded proteins for cell adherence, which are absent in genomes of other pathogenic members of the genus *Clostridium*. These include homologs of the adhesion protein Lap from the genus *Listeria*, the polar flagella from the genus *Aeromonas* and type IV pili from the genus *Neisseria* [48–50]. The genes *cheA/cheY*, *lisR/lisK* and *virR/virS* from the VF class regulation are virulence factors in *Listeria monocytogenes* and were also detected in the *Terrisporobacter* type strains [51–53]. Genes for the production of the toxins hemolysin and cytolysin involved in host cell lysis were detected in every *Terrisporobacter* strain. Furthermore, several virulence factors were detected associated to capsule formation involved in antiphagocytosis and immune evasion. For the verified *Terrisporobacter* pathogen *T. glycolicus* the *ddhA* gene was detected encoding an O-antigen essential for virulence in *Yersinia enterocolitica* [54]. Homologs of iron uptake proteins have been identified with similarity to proteins from the genus *Haemophilus* (hemB/L) and *Vibrio* (vctC), which matched with the incomplete heme biosynthesis encoded by all three genomes. Overall, the abundance and diversity of virulence factors detected in the three *Terrisporobacter* type strains underline the pathogenic potential harbored by this bacterial genus.

**Table 2.**
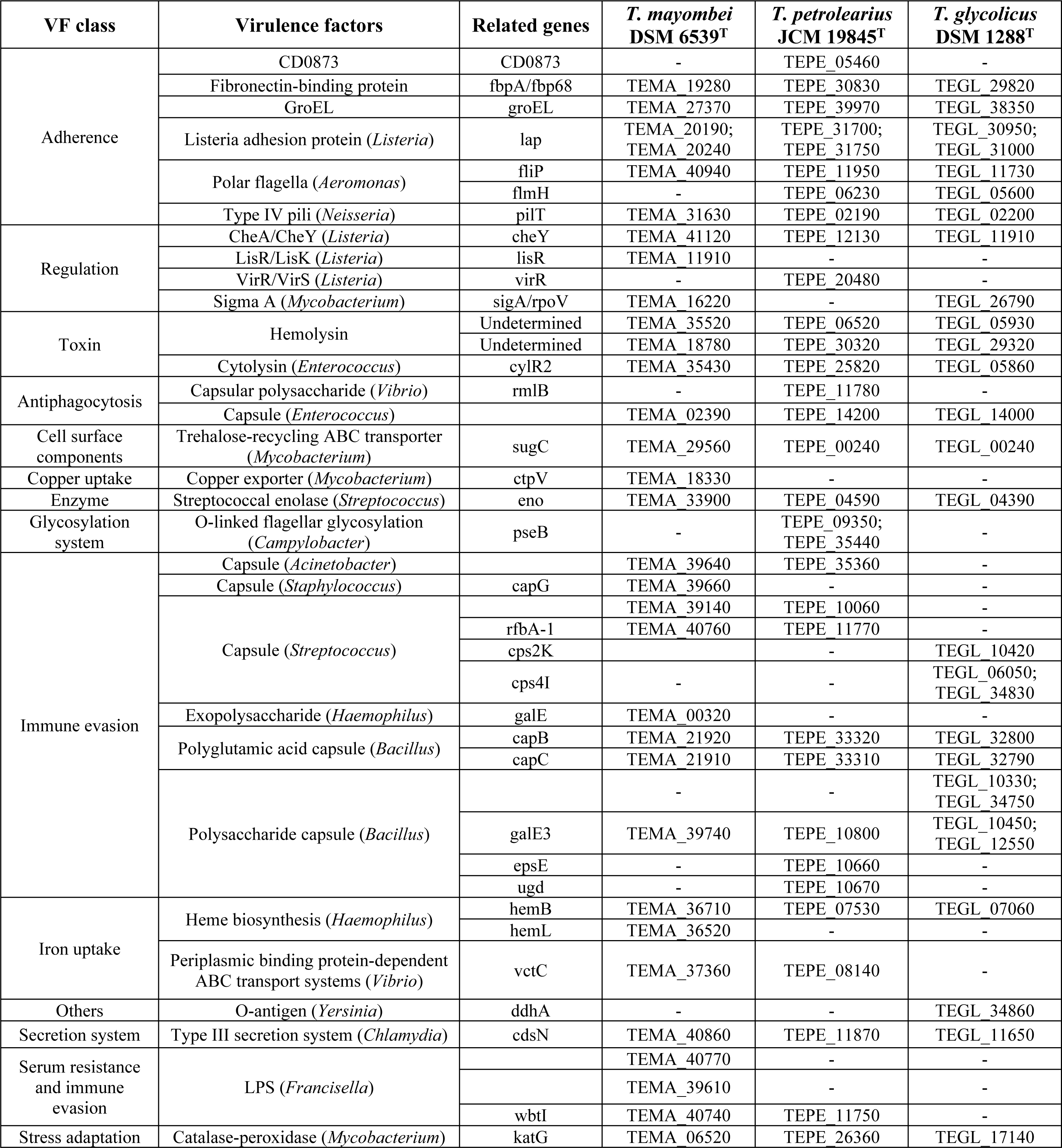
Virulence factor screen for three *Terrisporobacter* type strains with VFanalyzer using the VFDB.

### Phylogenomic analysis

The complete genome sequences were compared to available genome sequences of the *Terrisporobacter* genus by average nucleotide identity analysis based on a MUMmer alignment (ANIm) (Fig 7). The genomes of the type strains *T. glycolicus* DSM 1288^T^ and *T. mayombei* DSM 6539^T^ showed the highest identity to each other (93.5 %). This value was below the species delineation threshold of 95 % and supported the classification as separate *Terrisporobacter* species [55]. Using the same threshold, *T. glycolicus* DSM 1288^T^ and *T. petrolearius* JCM 19845^T^ also clustered as two distinct species with an identity of 93.1 %. However, the genome *T. petrolearius* JCM 19845^T^ showed the highest identity to the genome sequences of non-type *T. glycolicus* strains MGYG-HGUT-00005 (96.3 %), KPPR-9 (96.4 %), WW3900 (96.3 %), UBA8115 (95.8 %), FS03 (96.3 %), *T. petrolearius* UN03-225 (96.4 %) and *T. hibernicus* NCTC 14625^T^ (96.4 %). These identities were above the species identity threshold of 95 %, suggesting that these genomes belonged to the same species, namely *T. petrolearius*. Mitchell et al. reported a 67.4 % d4 dDDH value of the complete *T. hibernicus* type strain genome to the draft genome of *T. petrolearius* JCM 19845^T^ calculated by the Type Strain Genome Server (TYGS) [10,31]. TYGS submission of the complete genome of *T. petrolearius* JCM 19845^T^ yielded a d4 dDDH value of 69.2 % to the complete *T. hibernicus* genome. However, TYGS recommends the use of the dDDH values d0 and d6 for the comparison of complete genomes, as they also take the genome length into consideration in their calculations [56]. The dDDH values for d0 and d6 between the two complete genomes of *T. petrolearius* JCM 19845^T^ and *T. hibernicus* were 72.2% and 74.1 %, respectively. These dDDH values were above the generally applied species threshold of 70 % and thereby further supported the classification of the isolate *T. hibernicus* as member of the *T. petrolearius* species. Our phylogenomic analysis based on complete type strain genomes suggested the existence of subspecies clusters within the *T. petrolearius* species rather than two distinct species. With 99.6 % identity between the genome sequences of *T.* “*othiniensis*” 08-306576 and *T.* “*muris*” DSM 29186 both names describe isolates of the same species as already described by Afriza et al. [9]. Both isolates were not validly published under the ICNP as *T.* “*othiniensis*” was not deposited in a culture collection and *T.* “*muris*” is only deposited in one culture collection. Hence, no type strain and valid name description are currently available for these *Terrisporobacter* strains. In addition, the genome sequence of the isolate *T. mayombei* MSK4.1 showed an identity of 99.7 % to the genomes of *T.* “*othiniensis*” and *T.* “*muris*” and hence represents another isolate of this species. The genome sequence of *T. mayombei* ban5-GNNATD2-55 isolate from human feces was below the species delineation threshold of 95 % and thereby represented another undescribed *Terrisporobacter* species. The ANIm analysis of metagenomic assembled genomes from the *Terrisporobacter* genus indicated the existence of at least three more *Terrisporobacter* species without available isolates.

**Fig 7.**
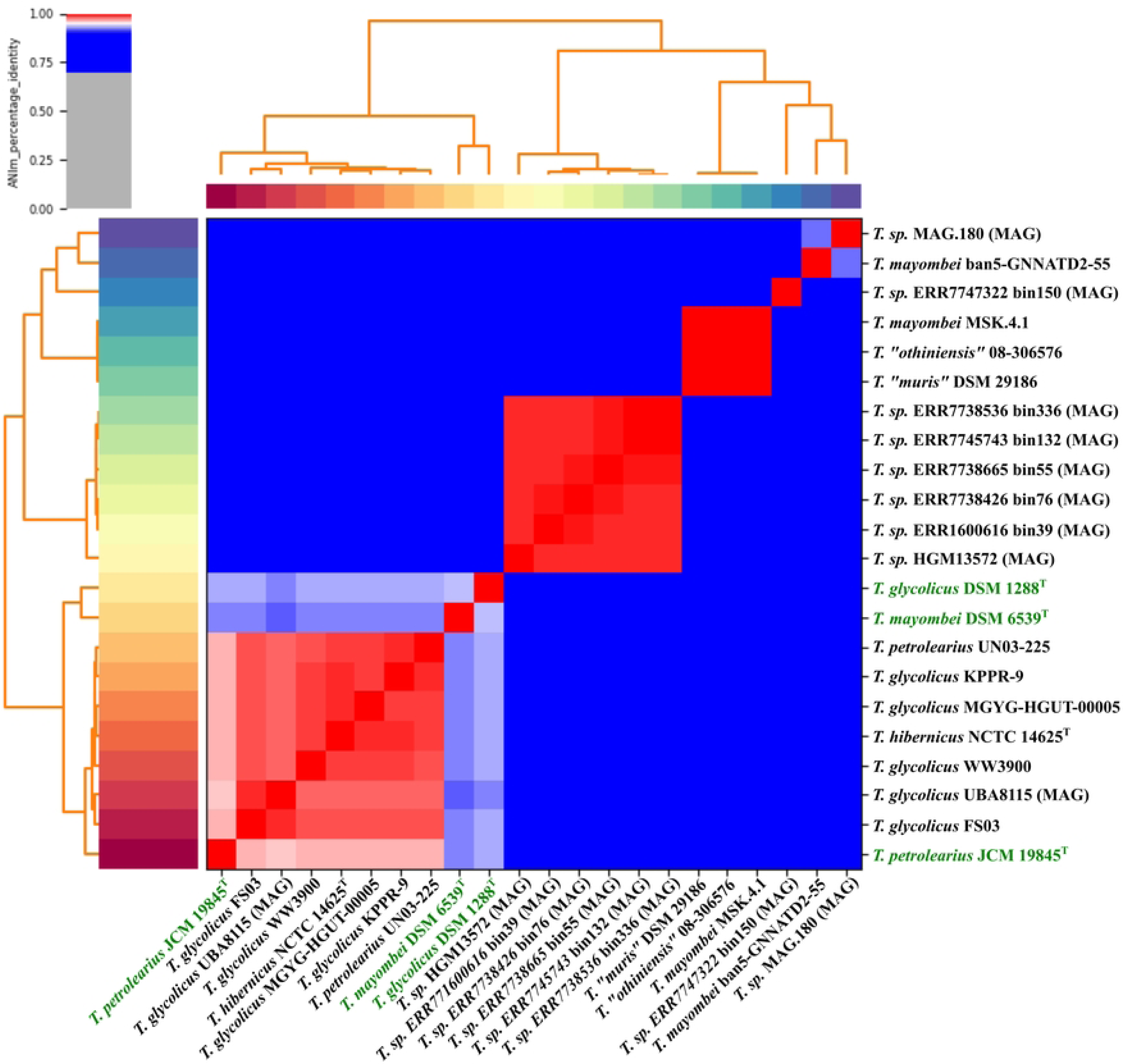
Phylogenomic analysis of *Terrisporobacter* genome sequences. The legend for sequence identities is shown in the top left corner. Sequence identities higher than 95 % are shown in red, while lower values are colored grey to blue. The three type strains of the genomes sequenced from the *Terrisporobacter* genus in this study are highlighted in green. Genome sequences derived from metagenomic assemblies are indicated by MAG in parentheses.

## Conclusions

Complete genomes for the three *Terrisporobacter* type strains of *T. mayombei*, *T. petrolearius* and *T. glycolicus* were obtained and revealed a *Terrisporobacter* characteristic 21 kb plasmid encoding a gene cluster for the reductive proline-dependent branch of the Stickland reaction. Functional annotations of the genomes showed species-specific auxotrophies for amino acids and vitamins. The genetic prerequisites for acetogenesis were given in all three *Terrisporobacter* type strains and the WLP could successfully be reconstructed. Our metabolic analysis supports future research for biotechnological applications of acetogenic strains from the genus *Terrisporobacter*. The detection of multiple potential virulence factors also demonstrated a pathogenic potential harbored by the genus *Terrisporobacter*. Phylogenomic analysis of available *Terrisporobacter* genomes suggested a reclassification of most isolates now termed *T. glycolicus* into *T. petrolearius*.

## Acknowledgements

We thank Melanie Heinemann for technical assistance and Avril von Hoyningen-Huene for proofreading.

## Supporting information

**S1 Table**. **Kits and program versions for Nanopore sequencing**.

**S2 Table**. **ANIm percentage identity and alignment length values.** ANIm percentage identity values (white, in %) and corresponding alignment lengths (grey, in bp) for available *Terrisporobacter* genomes at NCBI Genbank.

